# Sclerostin Deficiency Alters Peripheral B Lymphocyte Responses in Mice

**DOI:** 10.1101/357772

**Authors:** Arthur Chow, Jourdan Mason, Larrisha Coney, Jamila Bajwa, Cameron Carlisle, Anna Zaslavsky, Yvette Pellman, Marcos E. García-Ojeda, Aris Economides, Gabriela G. Loots, Jennifer O. Manilay

**Author notes:** **Corresponding Author:** Jennifer O. Manilay, PhD, University of California - Merced, School of Natural Sciences, 5200 North Lake Road, Merced, CA 95343.

## Abstract

Understanding how changes in bone physiology and homeostasis affect immune responses will inform how to retain strong immunity in patients with bone disease and in aged individuals. We previously identified sclerostin (*Sost*) as a mediator of cell communication between the skeletal and the immune system. Elevated bone mineral density in *Sost*-knockout (*Sost*^-/-^) mice contributes to an altered bone marrow microenvironment and adversely affects B cell development. B cells originate from hematopoietic stem cells within the bone marrow and mature in peripheral lymphoid organs to produce antibodies in response to infection and/or vaccination. In this study, we investigated whether the aberrant B cell development observed in the bone marrow of *Sost*^-/-^ mice extends to peripheral B cells in the spleen during immune challenge, and if these changes were age-dependent. Concomitant with more severe changes in bone architecture, B cell development in the bone marrow and in the spleen worsened with age in *Sost*^-/-^ mice. B cell responses to T-independent antigens were enhanced in young *Sost*^-/-^ mice, whereas responses to T-dependent antigens were impaired. Our results support the hypothesis that the adverse effects of B cell development in the *Sost*-deficient bone marrow microenvironment extends to the peripheral B cell immune response to protein antigens, and suggest that the B cell response to routine vaccinations should be monitored regularly in patients being treated with sclerostin antibody therapy. In addition, our results open the possibility that *Sost* regulates the T-independent B cell response, which might be applicable to the improvement of vaccines towards non-protein antigens.

## Introduction

Sclerostin-antibody therapy (Scl-Ab, romozosumab) has shown great promise for the treatment of osteoporosis, but the possible side effects of Scl-Ab administration and sclerostin depletion outside of the skeletal system are incompletely understood. Our previous work in the global *Sost*^-/-^ mouse revealed that B lymphocyte development in the bone marrow (BM) was adversely affected, resulting in reduced mature B cell numbers in the BM with high apoptotic activity ^(1)^ and our recent work using conditional *Sost*^-/-^ mice demonstrated a key role for *Sost* in mesenchymal stem cells on B cell maturation ^(2)^. Despite this reduction in the BM, no differences in B lymphocyte number were observed in the spleen. These results suggested that the obstruction in B cell development in *Sost*^-/-^mice was limited to bone marrow B cells, and that any B cells that completely matured then migrated to the spleen to fulfill normal immune functions in the periphery. In our original study, we focused our analyses on mice at peak bone mineral density (12-16 weeks of age), but patients enrolled in Scl-Ab clinical trials span an age range of 55-90 years ^(3,4)^. To more closely align our investigations with potential clinical outcomes of Scl-Ab administration in older individuals, we compared the effects of sclerostin deficiency on B cell development and the humoral immune response in young and aged mice.

## Materials and Methods

### Mice

*Sost^-/-^* mice have been previously described ^(1)^. Mice of both sexes were used on the C57BL/6 background and housed in sterile, microisolator cages with autoclaved feed and water. The UC Merced IACUC approved all animal work.

### Flow cytometric analysis of bone marrow and spleen cells

Procedures for euthanasia, dissection and preparation of femurs, tibias and spleens were performed as previously described ^(1)^. Details on bone marrow cell harvest and antibody staining are described in Supporting Information.

### Immunizations and ELISA

Pre-immune serum was collected from all mice. Mice were injected intraperitoneally with sterile PBS or NP_55_-Ficoll (25μg/100μl PBS) or NP_15_-OVA (50μg/100μl PBS), (Biosearch Technologies, Inc.). Mice treated with NP-OVA received two injections; the first in the presence of Alhydrogel adjuvant (InvivoGen, Inc.) and the second 14 days later without adjuvant. Serum was collected 8 days after NP-Ficoll injection or 7 days after the second NP-OVA injection, and was used immediately or stored at −80°C. ELISA plates (Fisherbrand) were plated with serial dilutions of purified mouse IgM, IgG3 or IgG1 (BioLegend) to create a standard curve. Sample wells were coated with 1μg of NP-BSA (Biosearch Technologies) in 100μl of 0.1M Na_2_PO_4_ pH 9.0 for 12-18 hours at 4°C, rinsed with (PBS-0.05% Tween 20) twice and the plate blotted to remove excess liquid. Blocking buffer (1.0% BSA in PBS-0.05% Tween 20, 0.05% sodium azide) was then incubated for 1 hour at room temperature, and wells washed and blotted. Diluted serum samples (100μl) were plated into NP-BSA coated wells, incubated overnight at 4°C, and wells washed and blotted 3 times. Horseradish peroxidase (HRP)- conjugated anti-mouse IgM, IgG3 or IgG1 (Southern Biotech, Inc.) was incubated for 30 minutes at room temperature, and wells washed and blotted 4 times. TMB Substrate Solution (Vector Laboratories, Inc.) was prepared per the manufacturer’s instructions, added to each well, and incubated for 20 minutes at room temperature. Fifty microliters of Stop Solution (1N sulfuric acid) was added to each well, and the absorbance read at 450 nm on a plate reader (Perkin Elmer) within 30 minutes. Serum concentrations of NP-specific IgM, IgG3 and IgG1 were calculated using the standard curve.

### Statistics

All data were expressed as the mean ± standard deviation. For flow cytometry results, statistical analysis was done using Student’s t-test with a two-tailed distribution, with two-sample equal variance (homoscedastic test). For ELISA, both t-tests and ordinary one-way ANOVA with multiple comparisons tests were performed. For all tests, p< 0.05 was considered to be statistically significant.

## Results

### Sost^-/-^ mice display age-related changes in B cell populations in the bone marrow and spleen

Changes in bone architecture are evident as early as 5 weeks of age in *Sost^-/-^* mice ^(5)^. To determine whether *Sost^-/-^* mice exhibit an age-dependent functional defect in B lymphocytes, we compared the B cell frequencies and numbers in the BM and spleens of “young” (1-3 months of age) and “aged” (>11 months of age) *Sost^-/-^* mice. We found that young *Sost^-/-^* mice displayed similar total B cell (B220^+^ IgM^+^ and CD19^+^ B220^+^) frequencies and absolute cell counts in the BM to that of age-matched wild-type (WT) controls (Figure 1A-C and Supp. Figure 1). In addition, the frequency of mature B220^high^ IgM^+^ B cells in the young *Sost^-/-^* BM was decreased (Figure 1G), but this did not result in a difference in cell number (Figure 1H). Aged *Sost^-/-^* mice displayed a more dramatic bone mineral density defect, where the bone marrow cavity was almost completely occluded. As expected, total BM cellularity (Figure 1A) and the frequency of B cells present in the BM were significantly decreased in these mice compared to age-matched controls (Figure 1A, B, G, H). We previously reported that in 12-16 week old *Sost^-/-^* mice, the block in B cell development begins at the pro-B cell stage and is maintained to Fraction D (B220^+^ CD43^+^ IgM^-^ IgD^-;^ late pre-B cell stage), and Fraction F (B220^+^ CD43^-^ IgM^+^ IgD^+^) was also reduced in the bone marrow in frequency and absolute number ^(1)^ In the current study, we did not observe any differences between the frequencies of any of the Hardy subsets ^(6)^ in young (< 3 months) control and *Sost^-/-^* mice (Figure 1I), but still observed a clear reduction in the absolute number of Fraction D, and increase in Fraction E, and reduction in Fraction F cell numbers in the *Sost^-/-^* mice (Table 1). The reduction in the absolute numbers of Fraction D and Fraction F cells was also observed in 8 week old mice treated with sclerostin-depleting antibody for 6 weeks (data not shown). In aged *Sost^-/-^* mice (>11 months), no difference between the frequencies of early pre-B (B220^+^ cKit^-^ BP1^+^ CD25^+^) and pro-B cells (B220^+^ cKit^+^ BP1^-^CD25^-^) ^(7)^ was observed, but there was an increase in the frequencies of Fraction D and Fraction E cells, and a reduced frequency of Fraction F cells (Figure 1J). This suggests that there is a progressive block in bone marrow B cell development beginning at the pro-B cells stage that becomes more severe with age in *Sost^-/-^* mice, which hinders the maturation of Fraction D and Fraction E cells to the Fraction F stage. Consistent with the reduced bone marrow cellularity observed in *Sost^-/-^* mice with age, all B cell fractions were significantly reduced in aged *Sost^-/-^* mice (Table 1).

**Figure 1.**
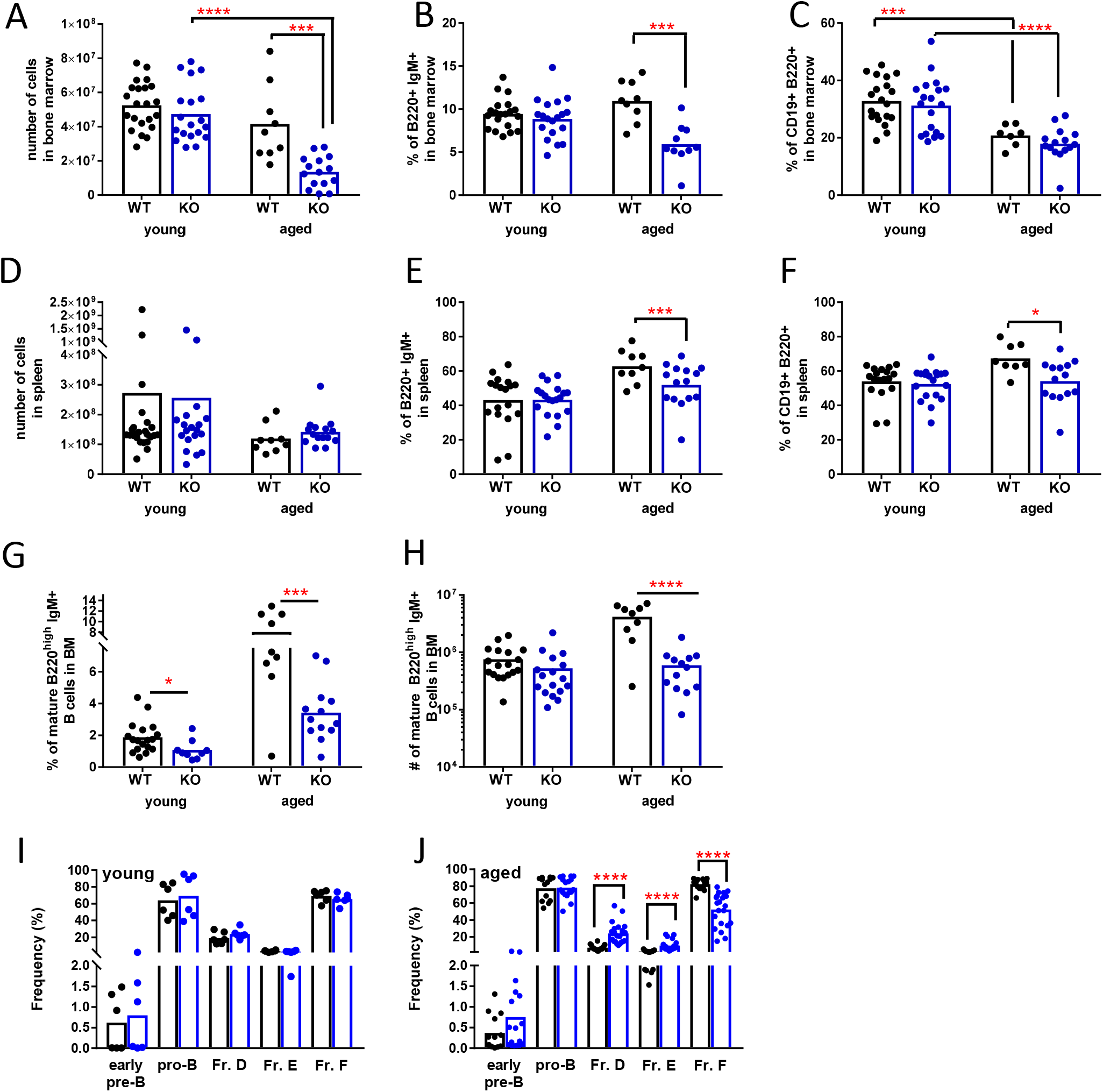
Comparison of B cell development in Sost^-/-^ mice by age. Total cellularity of BM (A) and spleen (D) in young (1-3 months) and aged (>11 months) WT and *Sost^-/-^* mice; frequency (%) of B220^+^ IgM^+^ and CD19^+^ B220^+^ cells in BM (B, C), and spleen (E, F), % and number of mature (recirculating) B220^high^, IgM^+^ (G, H) cells in bone marrow, and % of cells at distinct stages of maturation in the BM (I, J) of young and older WT and *Sost^-/-^* mice, based on flow cytometric analysis. Asterisks indicate statistical significance: ^*^p<0.05, ^***^p<0.001, ^****^p<0.0001. Each circle represents an individual mouse.

**Table 1.** Absolute Cell Numbers of B cell Bone Marrow Fractions.

| Table 1. Absolute Cell Numbers of B cell Bone Marrow Fractions |  |  |  |  |
| --- | --- | --- | --- | --- |
| Population<br>Mean ± SD | WT young | *Sost*<sup>-/-</sup> young | WT aged | *Sost*<sup>-/-</sup> aged |
| Early pre-B<br>(B220<sup>+</sup> cKit<sup>-</sup> BP1<sup>+</sup><br>CD25<sup>+</sup>) | 8.86 ± 9.62x10<sup>4</sup><br>n=6 | 8.66 ± 9.85x10<sup>4</sup><br>n=6 | 3.39 ± 3.27 x10<sup>4</sup><br>n=13 | **6.20 ± 6.08 x10<sup>3</sup> \*\*\***<br>n=17 |
| Pro-B (B220<sup>+</sup><br>cKit<sup>+</sup> BP1<sup>-</sup>CD25<sup>-</sup>) | 10.2 ± 5.56 x10<sup>4</sup><br>n=6 | 10.5 ± 7.04 x10<sup>4</sup><br>n=6 | 15.3 ± 9.90 x10<sup>4</sup><br>n=13 | **5.61 ± 5.16 x10<sup>4</sup> \*\*\***<br>n=17 |
| Fraction D (B220<sup>+</sup><br>CD43<sup>+</sup> IgM<sup>-</sup> IgD<sup>-</sup>) | 9.15 ± 2.03 x10<sup>5</sup><br>n=6 | **6.02 ± 2.63 x10<sup>5</sup> \***<br>n=6 | 3.82 ± 2.29 x10<sup>5</sup><br>n=18 | **1.21 ± 7.18 x10<sup>4</sup> \*\*\***<br>n=22 |
| Fraction E (B220<sup>+</sup><br>CD43<sup>-</sup> IgM<sup>+</sup> IgD<sup>-</sup>) | 1.96 ± 8.61 x10<sup>4</sup><br>n=6 | **7.04 ± 2.78 x10<sup>4</sup> \*\***<br>n=6 | 1.51 ± 8.04 x10<sup>4</sup><br>n=18 | **4.79 ± 3.12 x10<sup>4</sup> \*\*\***<br>n=22 |
| Fraction F (B220<sup>+</sup><br>CD43<sup>-</sup> IgM<sup>+</sup> IgD<sup>+</sup>) | 3.81 ± 1.65 x10<sup>6</sup><br>n=6 | **1.66 ± 0.66 x10<sup>5</sup> \***<br>n=6 | 5.05 ± 2.77 x10<sup>6</sup><br>n=18 | **4.36 ± 4.23 x10<sup>5</sup> \*\*\***<br>n=22 |
| *p<0.05, **p<0.01, ***p<0.001 |  |  |  |  |

After completing B cell development in the BM, B cells migrate to peripheral lymphoid organs, such as the spleen. Young *Sost^-/-^* and control mice displayed similar splenic B cell frequencies and numbers (Figure 1D-F). In contrast, aged *Sost^-/-^* mice contained a lower percentage of peripheral B lymphocytes in the spleens (Figure 1E-F). This was particularly surprising, as *Sost* expression has not been reported in the murine spleen ^(8)^.

### T-cell independent B cell responses are enhanced in young sclerostin-knockout mice

T-independent B cell responses are mediated by marginal zone (MZ) B cells in the spleen ^(9)^. We quantified the frequencies and absolute numbers of splenic marginal zone B cells in naive young and aged WT and *Sost^-/-^* mice using flow cytometry ^(10)^. No differences in the frequencies of MZ B cells between young and aged mice within the same genotype (e.g. young WT vs. aged WT) or different genotypes (young WT vs. young *Sost^-/-^*) were observed (Supp. Table 1). However, aged WT and *Sost^-/-^* mice expressed a lower absolute number of marginal zone B cells compared to their younger counterparts (although this was only statistically significant for the WT mice (Supp. Table 1). Therefore, it appears that MZ B cell numbers are reduced with aging, but not necessarily affected by the lack of *Sost*.

Peripheral B cell responses against T-independent antigens were tested using immunization with NP-Ficoll, a classic multimeric, repeating epitope-bearing T-independent type 2 (T1-2) antigen that simulates an immune response to polysaccharide antigens similar to those found on bacterial cell walls ^(11,12)^ (Figure 2A). We quantified the NP-specific T-independent immune response in *Sost^-/-^* mice by ELISA for NP-specific antibodies of the IgM and IgG3 isotypes in the peripheral blood ^(13)^ and compared these responses to age matched controls. Antibody titers in immunized mice were compared to pre-immunization levels, as well as to WT and *Sost^-/-^* mice that received injections of phosphate-buffered saline (PBS). Young immunized WT and *Sost^-/-^* mice both elicited a strong NP-specific response compared to saline-treated controls (Figure 2B, 2C). No differences in the IgM titers were observed in NP-Ficoll immunized young or aged *Sost^-/-^* mice compared to controls (Figure 2B, 2D). The levels of NP-specific IgG3 were slightly higher in the immunized young *Sost^-/-^* mice compared to age-matched immunized controls (Figure 2C). This IgG3 response in the immunized aged control and *Sost^-/-^*mice (Figure 2E) was similar, suggesting that aged *Sost^-/-^* mice have normal immunity to T-independent antigens.

**Figure 2.**
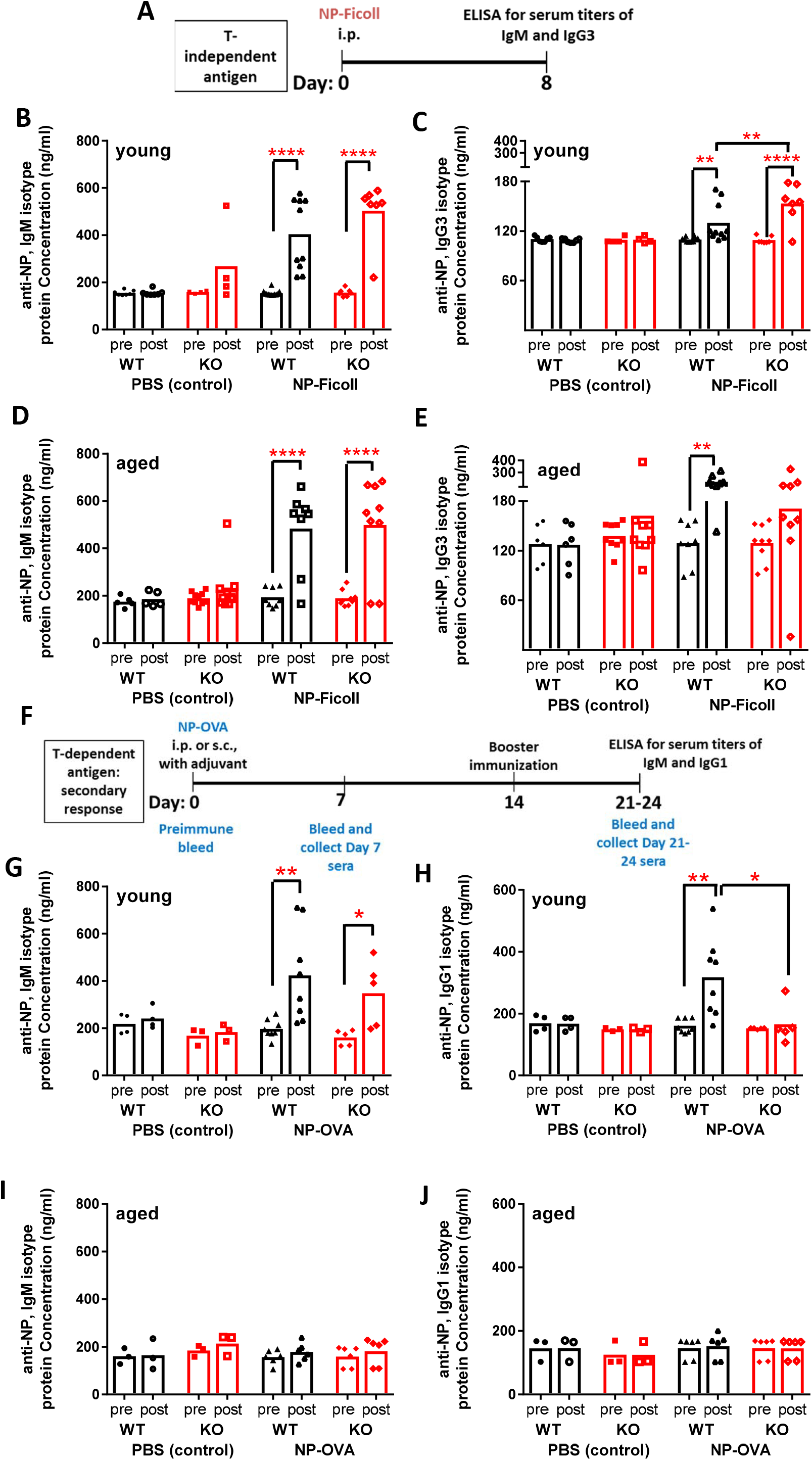
Humoral immune responses after vaccination in young and aged Sost^-/-^ mice. A-E) Analysis of T-independent B cell responses: Experimental design (A); Analysis of NP-specific IgM titers and IgG3 titers in young WT and *Sost^-/-^* mice (B, C), Analysis of NP-specific IgM titers and IgG3 titers in aged WT and *Sost^-/-^* mice (D, E). F-J) Analysis of T-dependent B cell responses: Experimental design (F); Analysis of NP-specific IgM titers and IgG1 titers in young WT and *Sost^-/-^* mice (G, H), Analysis of NP-specific IgM titers and IgG1 titers in aged WT and *Sost^-/-^* mice (I, J). Asterisks indicate statistical significance: ^*^p<0.05, ^***^p<0.001, ^****^p<0.0001. Each symbol represents an individual mouse.

### T-cell dependent B cell responses are diminished in young sclerostin-knockout mice

T-cell dependent B cell responses are mediated by follicular B cells ^(9,14)^. Frequencies and absolute number of follicular B cells were not affected by age or absence of Sost (Supp. Table 1). To determine whether *Sost^-/-^*mice respond to T-dependent antigens, *Sost^-/-^*and control mice were immunized with NP-OVA and B-cell responses were quantified. In this model, NP-OVA stimulates a B cell response to a protein antigen and requires induction of cytokines from CD4^+^ T helper cells to initiate the appropriate signals for isotype class switching from IgM to IgG1 ^(15,16)^ after a “booster” injection in B lymphocytes (Figure 2F). Young *Sost^-/-^* mice and controls mounted a similar IgM response to NP-OVA (Figure 2G), mirroring the response observed with the NP-Ficoll injections (Figure 2B). However, secondary immunization did not initiate isotype class switching to IgG1 in young *Sost^-/-^* mice (Figure 2H), compared to the clear presence of NP-specific IgG1 antibodies in the young WT controls. No significant differences in immune responses were observed between aged *Sost^-/-^*and WT controls (Figure 2I, 2J). These data suggest that *Sost*-deficiency might affect helper T cell maturation and functional response after stimulation. However, no differences in CD4 and CD8 T cell development in the thymus and spleen were evident in *Sost^-/-^* mice ^(1)^, and the ability of T cells from *Sost^-/-^* mice to proliferate after *in vitro* stimulation with concanavalin A was intact (data not shown), demonstrating that signal transduction downstream of the T cell antigen receptor is functional. However, subtle changes resulting in incomplete B cell activation are evident, as *in vitro* stimulation of *Sost^-/-^* B cells with anti-CD40 resulted in weaker blast formation ^(17)^ (Supp. Figure 2).

## Discussion

Recent reports on the clinical trials of Scl-Ab aimed at treatment for bone fragility diseases such as osteoporosis demonstrate promising results ^(3,4)^. Scl-Ab may also play a therapeutic role in many sclerostin-mediated diseases other than osteoporosis, such as rheumatoid arthritis ^(18)^, hypophosphatasia ^(19)^, osteogenesis imperfecta ^(20)^ and bone rebuilding in multiple myeloma ^(21)^. However, the impacts of Scl-Ab therapy on the immune system have not yet been examined. Our results show that the B cell response to protein antigens is diminished in younger *Sost^-/-^* mice, and these data may be relevant to human pediatric patients that receive Scl-Ab for treatment of osteogenesis imperfecta ^(22)^. On the other hand, the B cell response to protein antigens was universally weak in both older WT and *Sost^-/-^* mice, which might explain why relatively few of the adult patients receiving Scl-Ab (which itself is a protein antigen) mount an immune response to the Scl-Ab ^(3,4)^. That is, inhibition of the T-dependent antigen response by Scl-Ab may promote the efficacy and effectiveness of Scl-Ab therapy itself. We have not observed differences in the frequencies, numbers and activation capacity of T cells in *Sost^-/-^* mice, but further investigation is necessary to reveal if subtle differences exist within T cell subsets within the *Sost^-/-^* microenvironment, such as follicular T helper cells ^(23)^, which normally promote co-stimulation of B lymphocytes. Cells of the myeloid lineage can also promote cell responses by conventional B cells ^(24)^, providing an alternative area for experimentation.

Our previous studies demonstrated no effect on Sost deficiency on the *in vitro* B cell response to the T-independent type 1 (TI-1) antigen lipopolysaccharide ^(1)^. Thus, the enhanced ability of B cells in younger *Sost^-/-^* mice to respond to NP-Ficoll (a TI-2 antigen) *in vivo*, is intriguing. Responses to T-independent antigens can be mediated by innate-like B cells, such as unconventional B-1 lymphocytes (a fetally-derived cell population that persists during adulthood in the peritoneal cavity and mucosa, and may also develop postnatally in the BM ^(25,26)^), and splenic marginal zone B cells ^(27)^. Further investigation is required to determine the mechanisms underlying the enhanced IgG3 response to T-independent antigens in the younger *Sost*^-/-^ mice, such as a simple decrease in conventional B lymphocytes, and if sclerostin deficiency promotes expansion, activation and/or changes in Fc receptor expression in innate-like B cell subsets ^(28)^. There is evidence that the function of marginal zone B cells declines with age ^(29)^. The IgG3 response to T-independent antigens was somewhat higher in aged WT mice compared to younger WT mice, yet similar between younger and aged *Sost^-/-^* mice. This suggests that there is a role of *Sost* on innate-like B cells or their microenvironments ^(30)^ that may control the intensity of their T1-2 response to a basal (young) level.

The mechanisms that drive the relationship between Sost, Wnt and B cell development are not clearly understood. The role of Wnt signaling in B cell development is somewhat controversial. No differences in Wnt target gene expression was observed in B cells in the bone marrow of *Sost^-/-^* mice ^(1)^, suggesting that Wnt signaling in B cells is not driving the functional differences we have observed in the current study. In line with this, studies of B cell development in beta-catenin knockout mice demonstrated that B cell development is not dependent on canonical Wnt signaling; however, enhanced IgG3 class switching was observed in vitro, but no differences in T-independent or T-dependent B cell responses to antigen were observed in vivo ^(31)^. Mice that lack the Fanconi anemia gene *Fancc* experience progressive bone marrow failure, and *Fancc^-/-^* B cells display enhanced Wnt signaling, impaired B cell survival and impaired generation of antibody-secreting cells ^(32)^. The role of non-canonical Wnt signaling in B cell development has also produced conflicting results. Wnt5a-expressing stromal cells enhance the production and proliferation of B cell progenitors in vitro ^(33)^, but studies of Wnt5a haploinsufficiency in vivo showed that Wnt5a inhibits proliferation of B cell progenitors ^(34)^ and Wnt5a overexpression seemed to favor myeloid cell development over lymphoid development ^(35)^. Further investigation is required to settle these controversies and then to identify the specific roles of Sost in the communication between the skeletal and immune systems.

In conclusion, we demonstrate that sclerostin deficiency affects the peripheral B lymphocyte immune response to foreign antigens in mice, which could be relevant to the ongoing human clinical trials of Scl-Ab therapies. Our data suggest that B cell responses to both T-independent and T-dependent antigens, such as those found in routinely administered vaccines, be regularly monitored in patients that receive Scl-Ab treatment, as these therapies might inadvertently result in impaired B cell responses.

## Acknowledgments

This work was supported by University of California, Merced faculty research funding to J.O.M. and a student research grant to A.C. GGL works under the auspices of the U.S. Department of Energy by Lawrence Livermore National Laboratory under Contract DE-AC52-07NA27344. Authors’ roles: AE and GL provided the *Sost^-/-^* mice to begin our colony at UC Merced. Study design: JOM. Study conduct: JOM. Data collection: AC, JM, LC, JB, CC, AZ, YP, MGO. Data analysis: AC, JM, LC, JB, CC, AZ, YP and JOM. Data interpretation: JOM and GL. Drafting manuscript: JOM. Revising manuscript content: GL, MGO and JOM. Approving final version of manuscript: JOM. JOM takes responsibility for the integrity of the data analysis. The authors thank the staff of the Department of Animal Research Services and the Flow Cytometry Core of the Stem Cell Instrumentation Foundry at UC Merced for excellent animal care and technical support.

